# Sequence preparation is not always associated with a reaction time cost

**DOI:** 10.1101/2025.11.11.687917

**Authors:** Armin Panjehpour, Mehrdad Kashefi, Jörn Diedrichsen, J. Andrew Pruszynski

## Abstract

The extent to which a sequence of movements is prepared before initiating the first movement is a longstanding question in motor neuroscience. The observation that reaction time (RT) increases for longer sequences has been used as evidence of sequence preparation – reflecting the additional demands of preparing multiple movements before initiating a sequential action. However, many processes contribute to RT, making it unclear whether the observed RT increases specifically reflect sequence preparation. For example, with longer sequences, participants face greater ambiguity in selecting their first movement in the sequence. Here, we test how much of the observed RT increases can be explained by the first-target ambiguity when reaching toward spatial targets. In our paradigm, we independently manipulate: (i) the number of future targets displayed, (ii) the number of targets to be acquired, and (iii) the spatial arrangement of the targets. This approach allows us to vary the demands of sequence preparation and first-target ambiguity, thereby enabling a direct assessment of their respective influence on RT. We report that RT increases with additional sequence elements but that this effect is fully explained by the ambiguity in selecting the first reach target. That is, sequence preparation causes no RT increase. In fact, when first-target ambiguity is eliminated, RT is constant across the number of displayed targets even though kinematic analysis reveals that participants have prepared a sequence. Together, these results indicate that preparing multiple reaches to spatial targets does not impose additional temporal costs relative to preparing a single reach.

## Introduction

Many everyday actions, such as typing or driving, consist of sequences of movements that must be performed with both speed and accuracy^1^. Previous studies suggest that, to effectively carry out such actions, individuals prepare multiple future movements before initiating the sequence^2–11^.

A key observation often taken as support for the idea that individuals prepare multiple future movements before initiating the sequence is that reaction time (RT) — the time from the go-cue to the onset of the first movement — systematically increases with the number of sequence elements displayed to the participants, presumably reflecting the added demands of preparing future movements in advance. However, RT is an indirect measure composed of multiple processes, including stimulus identification, target selection, and motor preparation^12–16^, whether for the first movement or multiple future movements in the sequence. Therefore, it remains unclear how much of the RT increases specifically reflect sequence preparation. For example, as more sequence elements are displayed, participants not only gain the opportunity to prepare a longer sequence of movements, but may also face greater ambiguity in selecting the first movement target among multiple options (i.e. first-target ambiguity)^17^. Here we disentangle the effects of first-target ambiguity and sequence preparation on RT in the context of visually-guided reaching towards spatial targets.

To disentangle first-target ambiguity and sequence preparation effects on RT, we independently varied the number of displayed future targets, the number of instructed movements, and the spatial arrangement of the targets. By precisely measuring movement kinematics^18–20^, we provide clear behavioral evidence that participants do prepare beyond the first movement before initiating a sequence. Specifically, we show that first movement kinematics are systematically shaped by future movement goals (i.e., coarticulation) and that this influence emerges immediately at the onset of the first movement. Importantly, although RT increases with additional sequence elements, we show that this RT increase is fully explained by the first-target ambiguity and that sequence preparation causes no additional RT increase. In fact, when first-target ambiguity is eliminated, coarticulation is still present but RT is constant across the number of displayed targets.

## Methods

### Participants

Fifteen participants (6 female), with an average age of 25.5 years (SD = 2.7), completed the main experiment which took 30 minutes. Fourteen participants (5 female) with an average age of 26.5 years (SD = 2.9), completed a second 45-minute control experiment. Ten participants completed both experiments. Data from the main experiment was used in Fig.3, 4, and 5, and data from the control experiment was used in Fig.6. All participants were right-handed with an average handedness score of 80.8 (SD = 13.3) according to the Edinburgh Handedness Inventory.

All participants provided informed consent and reported no history of musculoskeletal, neurological, or psychiatric disorders. Participants were reimbursed $15 per hour after each session. All procedures were approved by the Human Ethics Research Board at Western University.

### Apparatus

Participants performed the experiments using an exoskeleton robot (KINARM^21^, Kingston, Canada). They were seated in a height-adjustable chair, with their arm supported by the exoskeleton (Fig.1A). Participants moved their arm in the horizontal plane to reach targets presented on a virtual display, which also blocked direct view of their arm and hand. Participants were given online feedback about their hand position using a red circle with a radius of 0.3cm on the screen projected above the tip of their index finger (Fig.1A, B). Reach kinematics were recorded at 1000 Hz. A photodiode captured the exact timing of when targets were displayed on the screen.

**Figure 1.**
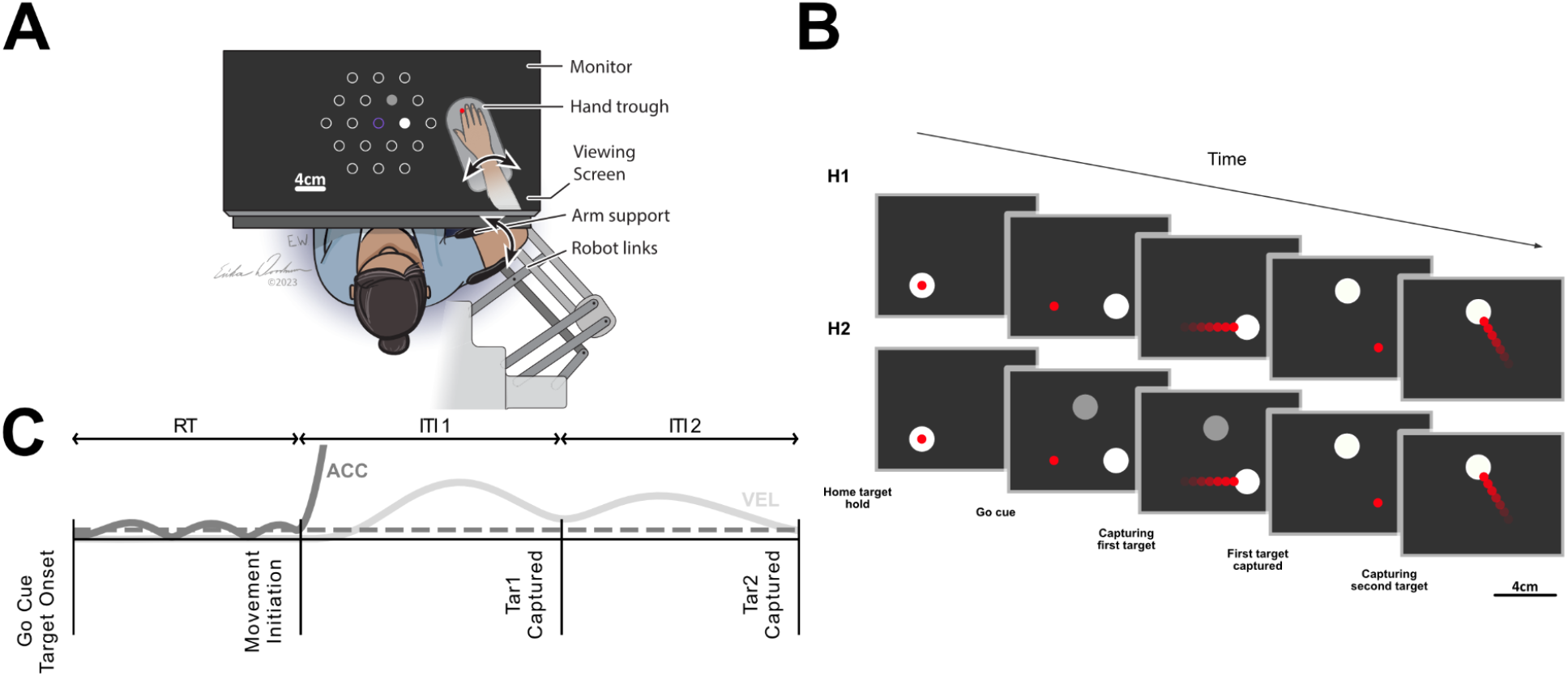
Apparatus and experiment design. A) Participants used the Kinarm exoskeleton robot to perform the reaches. Hand position was provided as feedback via a red circle, calibrated to the tip of the participant’s index finger. An example of a two-target sequence is shown; the full grid of potential targets was not visible to the participants. The order of the targets in the sequence was determined by their brightness. The home target is shown in purple here for visualization purposes only. B) Example sequence for the H1 and H2 horizon conditions. Participants start from the home target and are free to initiate their movement at the go-cue, which is also the target onset (Methods – Procedures). C) Acceleration and velocity profiles for an example trial (2-reach sequence – H2). RT was calculated based on the acceleration exceeding a specified threshold and staying above that threshold for 100 ms time points. Inter-Target-Intervals (ITIs) are the time between capturing each target. RT is excluded from ITI1.

### Procedures

In each trial, participants started from a fixed home target with a diameter of 1.5 cm positioned at the tip of their index finger when their shoulder and elbow joint angles were 45° and 90°, respectively. After a short delay, a series of four regularly paced auditory tones were played at 500 ms intervals. The last tone coincided with the disappearance of the home target and the appearance of the reach targets. Participants were free to initiate movement at any time after the last tone. The four auditory tones were provided to remove any ambiguity about the onset of the reach targets and the start of the trial. Reaching targets were generated from a two-layer hexagonal grid of potential targets with a diameter of 1.5cm and 4cm space between them (Fig.1A) and they were selected such that all reach distances were equal to 4 cm (i.e. each new target being one of the immediate neighboring targets to the current one).

In our main experiment, we manipulated three different factors independently (Fig.2). First, we manipulated the number of targets shown to the participant (Horizon), either one or two (H1, H2 - Fig.2B rows). In the H2 condition, the order of the targets was indicated by their brightness. Secondly, we instructed participants to perform either a single reach to the first target and ignore the second target if available, or perform a sequence of two reaches to capture two targets (Fig.2B). Finally, we manipulated the spatial arrangement of targets in the H2 condition (Fig.2B columns). In the non-spatially ordered condition, both targets were equidistant from the home position. In the spatially ordered condition, the two targets were positioned at different distances from the home position, with the first target being the closer one to the home target (Fig.2B columns).

**Figure 2.**
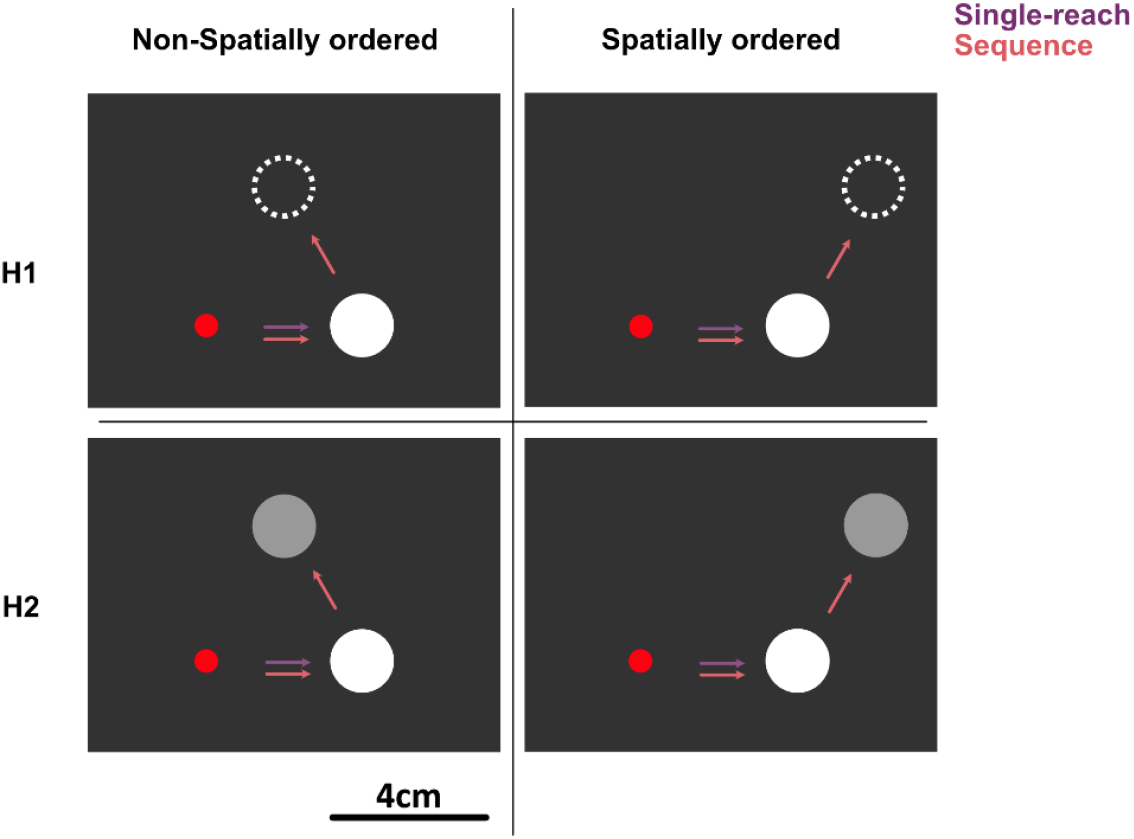
Experimental manipulations. The main experiment had three factors: horizon (H1, H2 - one or two targets visible on the screen), number of reaches (single reach or sequence of two reaches), and spatial arrangement of the targets (non-spatially ordered or spatially ordered). Snapshots at the go-cue are provided for this figure. All trials began from a fixed home target at the center. All reaching distances were 4cm, such that each new target was the immediate neighbor of the current one. Hand position was indicated by a red circle. Sequence order was indicated by the brightness of the targets in the H2 conditions. The dotted circles in the H1 condition indicate the second target that was not presented on the screen until the first target was captured. Arrows are shown for visualization purposes only.

During each trial, participants reached for targets in the instructed sequence, staying in each target for 50 ms to capture it (dwell time). If performing a sequence, the first target disappeared after it was captured, and the brightness of the second target was updated (Fig.1B).

The trial was terminated and participants were given error feedback if they committed any of the following actions: initiated movement before the fourth tone, exceeded the 800 ms time limit for each reach, failed to capture each target for the specified 50 ms dwell time, or directed their initial movement closer to a wrong first target. To calculate the initial movement direction, we used the velocity vector at the moment participants exited the home target. We then compared this direction with the positions of the visible targets on the screen. If the initial movement direction was oriented closer to the second target than the first target, the trial was terminated. This was done to discourage online corrections. The terminated trials were randomly repeated later in the session.

We also ran a single-session control experiment, using the same basic setup as the main experiment, to isolate the effect of spatial ordering on RT. This design avoided contamination from prior non-spatially ordered trials, since RT is known to be influenced by previous experience^14,16,22^. In the control experiment, participants only performed spatially ordered trials with horizon options of 1, 2, or 3 and longer sequences of 5 reaches. Targets were selected from a 3-layer hexagonal grid, and the first three targets were spatially ordered.

### Experiment Design

In the main experiment, participants completed two sets of 7 blocks. Each set included 7 blocks based on a 2×2×2 design (number of horizon × reaches × spatial arrangement), excluding one redundant condition: the single-reach, horizon-1 block was identical across spatial arrangements. Blocks were randomly ordered within each set. Each block consisted of 30 trials, totaling 420 trials for the session, which lasted approximately 30 minutes. Within each block, the number of reaches, the horizon, and the spatial arrangement of the targets remained consistent. The block instructions were displayed on the top left of the screen.

In the control experiment, participants completed 540 trials of horizon-randomized 5-reach spatially ordered sequences. This experiment lasted for 45 minutes.

### Time Analysis

RT was calculated based on the acceleration exceeding a threshold of 5*cm/s*^2^ and staying above that threshold for 100ms. We calculated the average of inter-target-intervals (ITIs) to quantify how fast each sequence is executed. Each ITI includes the time that the hand leaves a target and captures the next target (Fig.1C). Mean ITI for each condition in Fig.3A was calculated by first averaging ITIs within each trial for each participant, then taking the median ITI across all trials for each participant, and finally averaging those medians across all participants. Between-subject variability was removed in the RT and ITI plots by first subtracting each participant’s overall mean across conditions and then adding the mean across all participants, preserving the true average across participants.

### Decoding Analysis

We used the reach kinematics to decode the first and second targets at each time sample in two-reach sequence trials (Fig.3). For decoding the first target, there were 6 possible targets based on the grid (Fig.3B). For decoding the second target, we rotated all the kinematics and targets so that all of their first targets lie in the 0-degree direction, on the (4,0) coordinates. For each of the non-spatially ordered and spatially ordered conditions, there were two possible second targets after all the conditions were rotated (Fig.3C). We did 10 runs of 5-fold cross-validated multinomial logistic regression at each time sample for decoding the first and second targets independently. We used hand position, velocity, and acceleration in both horizontal and vertical axes together with a constant bias as the inputs to the decoder and scalar value labels as outputs. We ran the decoder for decoding each of the first and second targets, in both horizons, both spatial arrangements, and each participant separately. To calculate the chance level, we shuffled the labels and performed 5 runs of hold-out multinomial logistic regression. Decoding onset (Fig.3E) was defined as the first time point when decoding accuracy exceeded the chance level with 95% certainty and stayed above this threshold for at least 100 consecutive samples.

**Figure 3.**
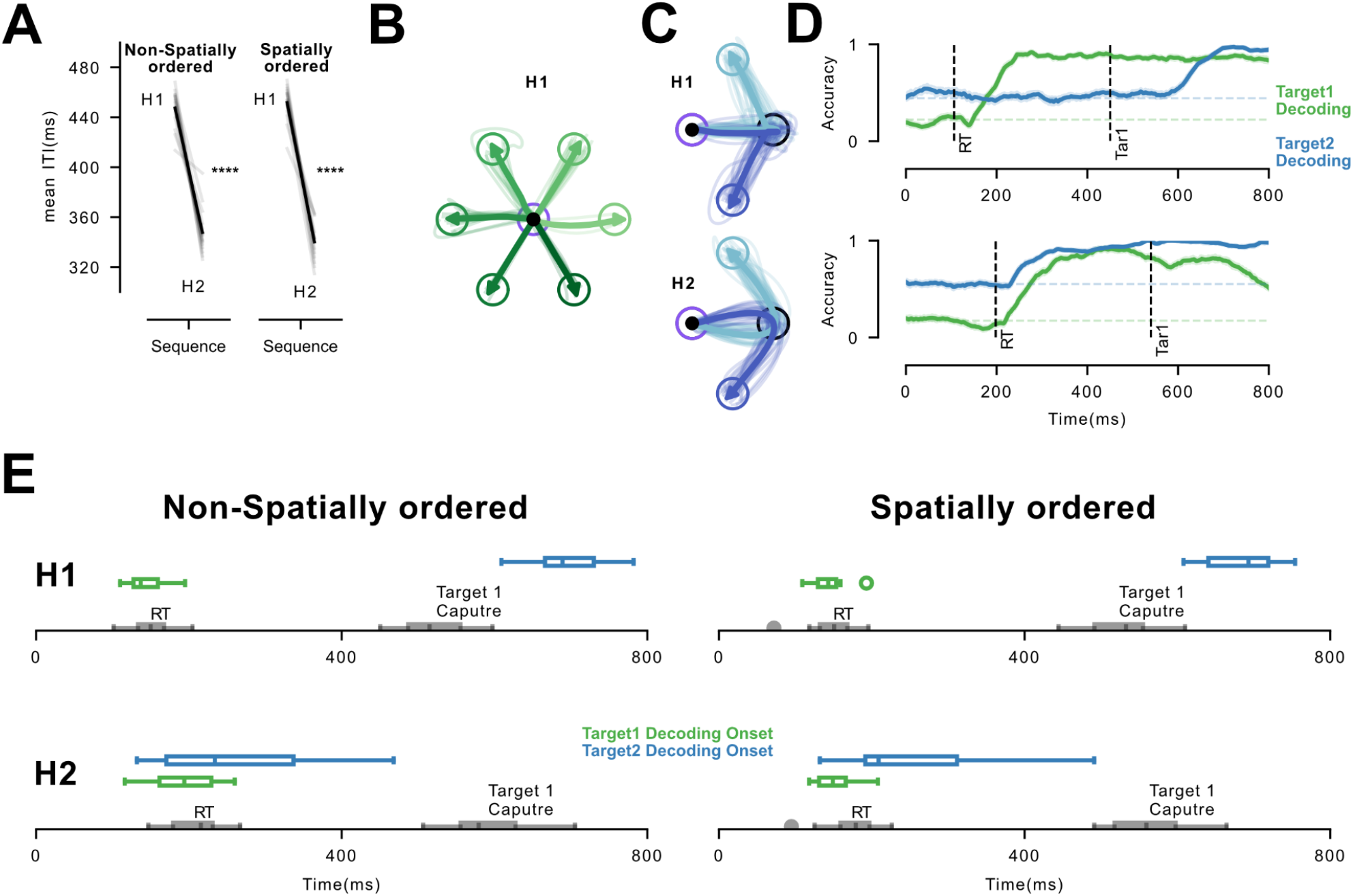
Preparation for multiple sequence movements before initiation. A) Inter-target-intervals (ITIs) for 2-reach sequences. The gray lines are the median over trials for single participants. The black lines are the average over the participants. **** signifies p-value < 0.0001 for individual t-tests across horizon levels. B) For one representative participant, green trajectories illustrate the first reach traces to all possible 1st targets in H1. C) For one representative participant, blue trajectories show the movement traces for the two sequences going to the same first target (black blank circle) but different second targets (dark and light blue) for H1 and H2 (C rows) for the non-spatially ordered trials. Thin lines are single-trial hand positions, and thick lines are the average over the trials. The home target is shown as a purple circle for visualization purposes only. D) For one representative participant, decoding accuracy (y-axis, 0 to 1) for predicting the 1st and 2nd targets is plotted over time for H1 and H2 (D rows). Vertical lines indicate RT and Target 1 Capture (Tar1). Solid lines represent cross-validated accuracy for the participant, and shaded areas indicate the s.e.m. across n runs. Decoding chance levels are indicated as horizontal dashed lines. E) Decoding onsets for the 1st and 2nd targets across all participants for all horizons and spatial arrangements for sequence trials based on kinematics. RT and Target 1 Capture are shown in gray for comparison. Box plots represent the median, first, and third quartiles of the data, and the whiskers extend from the box to the furthest data point lying within 1.5x of the inter-quartile range of the box. Dots represent any data points beyond the whiskers.

### Curvature Analysis

To calculate systematic curvature (Fig.4B) for each trial’s first reach, we first rotated all trials so that the first targets aligned at 0-degree direction. We then sampled 200 equally spaced points from each first reach trajectory. Next, we performed principal component analysis (PCA) on the matrix of y-axis values for the first reach across all trials, conditions, and participants (N trials × 200). Each trial was then projected onto the first PC, yielding a single signed value per trial. The absolute value of this scalar represents the curvature amplitude, while its sign indicates curvature direction. Finally, we multiplied this signed scalar by the sign of the rotated second target’s y-position to facilitate averaging across trials for different second targets, allowing us to detect systematic curvature patterns. Between-subject variability was removed when plotting curvature-value figures by first subtracting each participant’s overall mean across conditions and then adding the mean across all participants, preserving the true average across participants.

**Figure 4.**
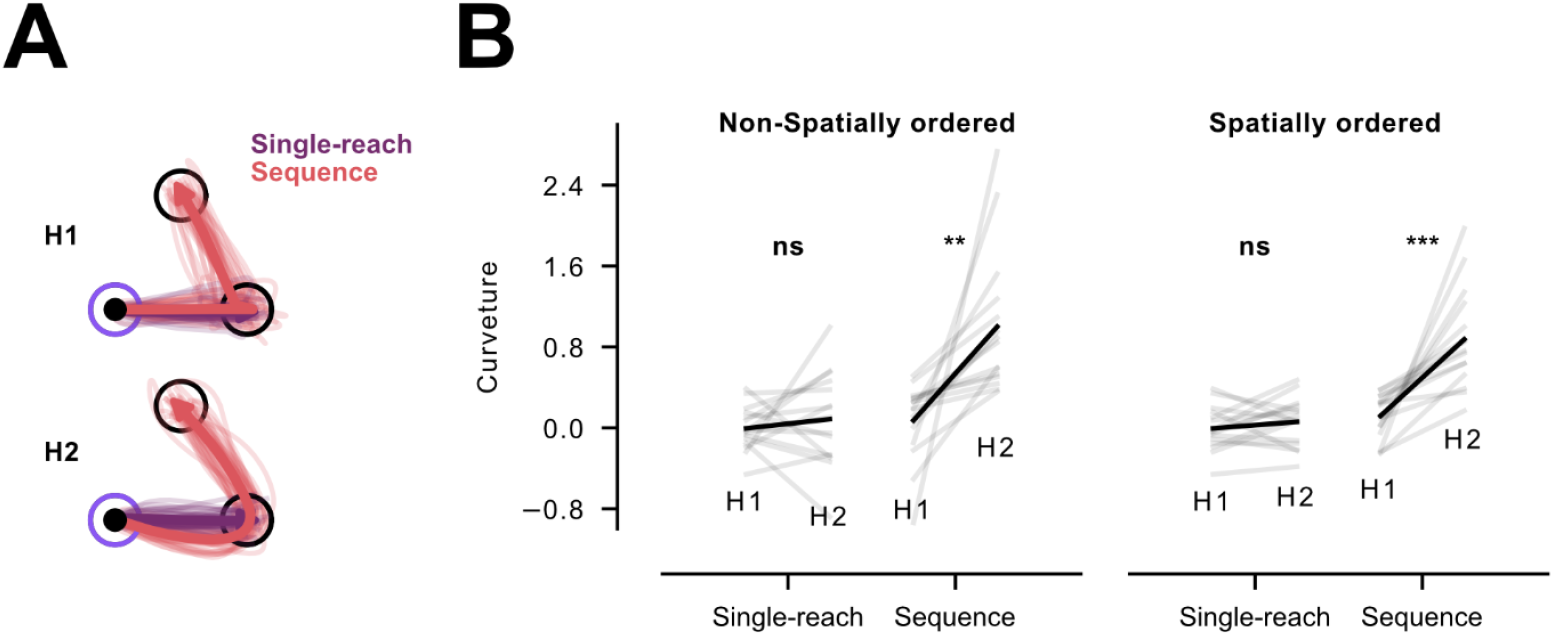
Coarticulation on the first reach in a sequence is a deliberate motor plan for the second target. A) For one representative participant, comparison of the first reaches of non-spatially ordered trials between single-reach and sequence conditions in H1 and H2. Thin lines are single-trial movement traces, and the thick lines are the average over all trials. Single-reach trials are indicated by purple and sequence trials by pink. The home target location is shown by the purple circle only for visualization purposes. Reach targets are shown by the blank black circles. B) First reach curvature values for all 8 conditions. The gray lines are the median over trials for single participants. The black lines are the average over the participants. Ns, **, and *** signify p-value > 0.05, p-value < 0.01, and p-value < 0.001 for individual t-tests across horizon levels.

### Statistical Analysis

We conducted three-way repeated measures ANOVA for all the curvature and RT analyses, incorporating the three experimental manipulations. The factors included spatial arrangement (2 levels), horizon (2 levels), number of reaches (2 levels), and their interactions. We also conducted two-way repeated measures ANOVA for the ITI analysis, incorporating spatial arrangement and horizon as the factors. We conducted paired t-tests to compare the horizon factor levels within each reach number and spatial arrangement condition. For the control experiment, we performed a one-way repeated measures ANOVA across the three levels of horizon. Detailed descriptions of the statistical tests are provided in the text accompanying the Results section.

## Results

Our main goal was to test whether more than one reach is prepared before initiating a sequence of reaches and to disentangle the first-target ambiguity effects from sequence preparation effects on RT. To do so, we manipulated three different factors in a reaching task. First, we manipulated the number of targets shown to the participant, either one or two (Horizon 1 - H1 or Horizon 2 - H2; Fig.1B & Fig.2 rows). This manipulation determined whether participants could use information about future targets to prepare beyond the first reach (Methods – Procedures). Second, we instructed participants to perform either a single reach to the first target and ignore the second target (if available) or perform a sequence of two reaches to both targets (Fig.2B). By keeping visual complexity (i.e. horizon) constant and manipulating the number of reaches, we dissociated sequence preparation effects on RT from the ambiguity in selecting the first reach target among the presented targets (i.e. first-target ambiguity). Third, we manipulated first-target ambiguity directly by varying the spatial arrangement of targets in the H2 condition (Fig.2B columns). In non-spatially ordered trials, both targets were equidistant from the home position, forcing participants to only rely on brightness cues to identify the first target (Methods – Procedures). In the spatially ordered condition, the two targets were positioned at different distances from the home position, allowing participants to use both distance and brightness cues to identify the first target (Fig.2B columns, Methods – Procedures).

### Multiple reaches are prepared before the initiation of the sequence

#### Advance information speeds up sequence production

We first tested if participants in the sequence condition were faster when they could see two future targets prior to movement onset. We quantified how long it took for a participant to complete both reaches by calculating the average inter-target-interval (ITI, time between target acquisitions, Methods – Time Analysis). ITIs were shorter in the H2 as compared to the H1 condition in both non-spatially ordered and spatially ordered trials (Fig.3A - significant horizon effect, F_(1, 14)_ = 195.95, p = 1.26e^-9^). This result indicates that participants used the information about the future targets, resulting in the faster production of the sequence – either during RT or as the first reach was unfolding^11,23,24^.

#### Kinematics analysis confirms preparation of multiple targets before movement initiation

To assess whether preparation for the second target happens before the movement onset, we examined how early the second target could be predicted from the kinematics of the first reach. Kinematics for sequences with the same first target and two different second targets for one representative participant are shown in Fig.3C. As illustrated, the first reach is systematically curved away from the second target when participants can see the second target (i.e. H2). To quantify this, we conducted a decoding analysis (Methods – Decoding Analysis) at each time sample to identify when the second target could be predicted based on the kinematics of each trial (Fig.3D). As expected, in the H1 condition, the second target could only be decoded once the reach toward it had started (i.e. after capturing the first target, Fig.3D, first row). In contrast, in the H2 condition, the second target could be decoded significantly earlier, almost immediately after movement initiation towards the first target, suggesting that information about the second target influenced motor preparation before movement start (Fig.3D, second row). This pattern was consistent across participants and for both non-spatially ordered and spatially ordered conditions (Fig.3E).

#### Curvature on the first reach represents a motor plan for the second target

Next, we investigated whether the observed curvature in the first reach was simply an involuntary visual response to the presence of the second target as a visual distractor, rather than a motor plan for the second target^25^. If the second target were just a distractor, participants should curve their reach away from the second target even in the single-reach condition. To address this, we compared the curvature values across horizons between the single-reach and sequence conditions (Fig.4A & Fig.4B). We found that participants co-articulated (Methods – Curvature Analysis) their first reach in the sequence condition (non-spatially ordered: t_(14)_ = 3.73, p = 2.3e^-3^, spatially ordered: t_(14)_ = 4.94, p = 2.1e^-4^) but not in the single-reach condition (non-spatially ordered: t_(14)_ = 0.71, p = 0.49, spatially ordered: t_(14)_ = 0.97, p = 0.35). This result indicates that the observed coarticulation results from a motor plan that includes information about the second target.

### Sequence preparation does not cause a RT cost

The above results show that sequence preparation happens before movement initiation. However, does this process influence RT? Comparing the H1 and H2 conditions for the non-spatially ordered condition (Fig.5), we found an increase in RT (non-spatially ordered: significant horizon effect, F_(1, 14)_ = 74.38, p = 5.65e^-7^), which can partially or fully be attributed to the ambiguity in selecting the first reach target, consistent with Hick’s law^17^. Importantly, when participants were instructed to perform two reaches, there was no additional RT cost compared to the single-reach condition. This was true for both non-spatially ordered and spatially ordered conditions (Fig.5, no interaction between horizon and number of reaches, F_(1, 14)_ = 1.58, p = 0.23). This suggests that none of the RT cost in the sequence condition arises from sequence preparation; rather, it is entirely due to resolving the first-target ambiguity.

**Figure 5.**
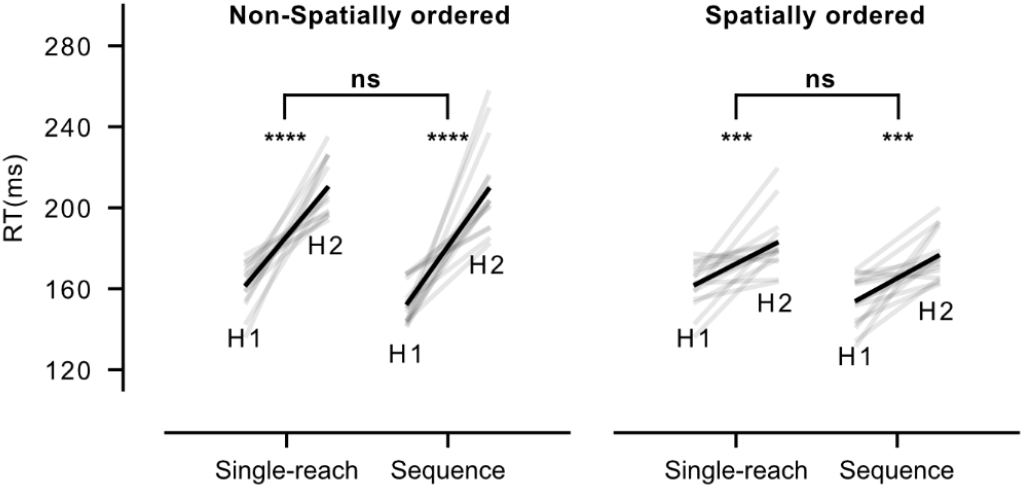
Sequence preparation does not cause any RT cost. RTs for all 8 conditions. The gray lines are the median over trials for single participants. The black lines are the average over the participants. *** and **** signify p-value < 0.001 and p-value < 0.0001 for individual t-tests across horizon levels. Ns signifies the non-significant interaction of horizon and number of reaches for ANOVA.

Next, we asked if the observed RT cost due to the first-target ambiguity could be reduced or eliminated through spatial ordering of the targets, which potentially makes resolving the ambiguity easier (Fig.2B right-column, Methods – Procedures). Indeed, the RT cost across horizons in the spatially ordered trials was significantly lower than in the non-spatially ordered trials (significant interaction between horizon and spatial arrangement, F_(1, 14)_ = 32.16, p = 1e^-4^), though it was not entirely eliminated (spatially ordered: significant horizon effect, F_(1, 14)_ = 50.15, p = 5e^-6^, Fig.5). This suggests that spatially ordering the targets makes resolving the first-target ambiguity easier.

#### Spatial ordering completely eliminates the first-target ambiguity

In the previous section, we found that the RT cost due to first-target ambiguity was significantly lower in spatially ordered trials compared to non-spatially ordered trials, although some small cost remained (Fig.5). However, RTs can be influenced by factors such as prior experience and the experimental context^14,16,22^. This led us to test whether the small but significant RT cost observed in the spatially-ordered trials was influenced by the carryover effects from non-spatially ordered trials in previous blocks (Methods – Experiment Design). To explore this, we analyzed RT across three horizons in a separate session (Methods – Procedures) collected from participants performing sequences of 5 reaches in which sequence order was always cued spatially. We found no RT cost across horizons (Fig.6), suggesting that spatial ordering can fully mitigate the first-target ambiguity (F_(2, 26)_ = 2.07, p = 0.15) and that sequence preparation happens without any RT cost (Fig.5).

**Figure 6.**
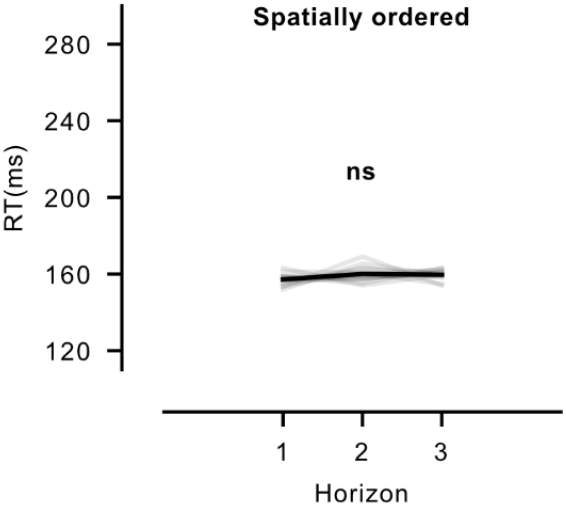
Spatial ordering fully mitigates first-target ambiguity, and sequence preparation happens without any RT cost. RTs for spatially ordered 5-reach sequence trials across 3 horizons in a separate session. The gray lines are the median over trials for single participants. The black line is the average over the participants. Ns signifies the non-significant horizon effect for ANOVA.

## Discussion

We found that participants do prepare beyond the first movement in a sequence of visually-guided reaches towards spatial targets but that this preparation imposes no additional RT cost compared to the preparation of a single reach. Instead, our findings indicate that RT increases with more sequence elements arise from the ambiguity associated with selecting the first movement target among several options^17^ (Fig.5). Our results align with recent findings showing that movement complexity and RT cost are largely independent^14^. To further probe the dissociation between first-target ambiguity and sequence preparation, we introduced a spatial arrangement condition in which the location of the targets, in addition to their brightness, served as an additional cue to resolve the first target. In this condition, the closer target was always the first reach target. We observed no RT cost across horizons, suggesting that spatial ordering can fully resolve the ambiguity in first reach target selection (Fig.6). First-target ambiguity effects on RT together with our coarticulation results, suggest that the preparation of multiple movements in target-directed reaching sequences occurs without incurring any additional RT cost compared to the preparation of a single movement (Fig.5 & Fig.6).

Additionally, unlike when spatially ordered trials were tested in a separate session, we found that mixing spatially-ordered trials with non-spatially ordered trials resulted in a smaller yet significant RT cost across horizon compared to non-spatially ordered trials (Fig.5). This likely reflects that RTs for spatially-ordered trials are biased by the presence of non-spatially ordered trials, in accordance with recent studies reporting habituation and contextual effects on RT^14,16,22^. Our results also align with recent evidence that movement preparation and initiation are independent processes^15^.

A previous study using finger-press sequences cued by digits on a screen has shown that RT increases as more sequence elements are displayed in advance^11^. The same paradigm has also shown that the first three presses in a sequence are executed more slowly when the time available for pre-planning is limited^26^. Our results now raise the possibility that a part of the RT cost observed in other paradigms such as finger sequences arises because of the need to resolve first-target ambiguity. However, it remains to be tested whether the absence of a RT cost for preparing multiple movements when reaching to spatial targets will generalize to other paradigms or effectors.

In contrast to reaching paradigms, finger movement paradigms often use abstract cues like digits and symbols, and thus, they are associated with a slower stimulus-response (S-R) mapping^11,27–30^. For example, single finger movements cued by digits typically require ∼400–700 ms to be initiated^11,29^, whereas single reaching movements to directly presented spatial targets can be initiated within ∼130–300ms^15,31–34^. Even more direct mappings, such as spatial cuing instead of symbolic cuing, do not eliminate the substantial RT differences between finger and reaching movements^27,28,30^. This complex S-R mapping in finger movements requires additional time, and it also suggests that the RT increase for finger sequence preparation is less likely due to additional demands at the level of motoric planning, but rather it arises at the level of more cognitive planning processes^35,36^. Given these differences in S–R mapping between reaching and finger movements, sequence preparation may impose an RT cost for finger-press sequences, even when first-target ambiguity is accounted for.

An important difference between the underlying mechanisms of reaching and other types of movements, such as finger-presses, is the rapid visuomotor pathways that link visual input to motor output for reaching movements^37–39^. These fast motor responses are potentially supported by the interplay of cortical and subcortical circuitries^40^. Within the dorsal stream^41^, the posterior parietal cortex (PPC) transforms visual target locations from retinal coordinates into hand- and body-centered reference frames^42^. At the cortical level, the primary motor cortex (M1) is a key component of fast feedback response circuits^43–52^, receiving inputs from multiple subcortical areas and sending outputs both to these subcortical regions^53,54^ and to the spinal cord^55^. Studies have also reported rapid responses in the forelimb area of M1 to visual stimuli^56–58^. At the subcortical level, the superior colliculus (SC), via the tecto-reticulo-spinal pathway, makes connections to arm muscles^59,60^ and shows modulations to movement-direction during both preparation and execution of single reaches^61,62^. Although these rapid visuomotor pathways are mostly studied in the context of single reaching movements, they may also support the rapid preparation and execution of reaching sequences. In fact, previous studies have shown sequence-specific activity in PPC^63^, as well as sequence-specific modulation of long-latency reflexes^19^. As for M1, a recent electrophysiological study employing a two-reach sequence, without co-articulation and with matched first reaches across different sequences, suggests that M1 does not participate in joint preparation of multiple reaches before sequence initiation^23^. However, the role of M1 in preparing multiple sequence elements when co-articulation is present from the start, as in the current study, remains unclear and will be the focus of future investigation.

## Author Contributions

Conceptualization: AP, MK, JD, JAP

Data collection: AP

Analysis and visualization: AP

Supervision: JD, JAP

Writing –original draft: AP

Writing–review and editing: AP, MK, JD, JAP

## Acknowledgements

This work was supported by a CIHR Project Grant to JD and JAP (PJT-175010). J.A.P received a salary award from the Canada Research Chairs program.

